# The mitochondrial deoxyguanosine kinase is required for female fertility in mice

**DOI:** 10.1101/2023.11.12.566728

**Authors:** Yake Gao, Rui Dong, Jiacong Yan, Huicheng Chen, Lei Sang, Xinyi Yao, Die Fan, Xin Wang, Xiaoyuan Zuo, Xu Zhang, Shengyu Yang, Ze Wu, Jianwei Sun

**Affiliations:** Center for Life Sciences, Yunnan Key Laboratory of Cell Metabolism and Diseases, State Key Laboratory for Conservation and Utilization of Bio-Resources in Yunnan, School of Life Sciences, Yunnan University, Kunming, China; Department of Reproductive Medicine, the First People’s Hospital of Yunnan Province, Kunming, China; NHC Key Laboratory of Preconception Health Birth in Western China.; Department of Cellular and Molecular Physiology, Penn State College of Medicine, Hershey, PA, USA

**Keywords:** DGUOK, infertility, mitochondria, nucleoside salvage pathway, oocyte maturation

## Abstract

Mitochondrial homeostasis plays a pivotal role in oocyte maturation and embryonic development. Deoxyguanosine kinase (DGUOK) is a nucleoside kinase that salvages purine nucleoside in the mitochondria and is critical for mitochondrial DNA replication and homeostasis in non-proliferating cells. *Dguok* loss-of-function mutations and deletions lead to hepatocerebral mitochondrial DNA deletion syndrome. However, its potential role in reproduction remains largely unknown. We found that *Dguok* knockout results in female infertility. Mechanistically, the deficiency of DGUOK hinders ovarian development and oocyte maturation. Moreover, *Dguok* deficiency in oocytes caused a significant reduction in mitochondrial DNA and abnormal mitochondrial dynamics, and impaired germinal vesicle breakdown. Only few DGUOK-deficient oocytes were able to extrude the first polar body during *in vitro* maturation, and these oocytes showed irregular chromosome arrangement and different spindle lengths. In addition, *Dguok* deficiency elevated reactive oxygen species and accelerated apoptosis in oocytes. Our findings reveal novel physiological roles for the mitochondrial nucleoside salvage pathway in oocyte maturation and implicate that DGUOK is a potential marker for the diagnosis of female infertility.

## Introduction

Infertility poses a global health challenge, impacting approximately 15% of couples worldwide(1). The process of successful reproduction involves the normal maturation of oocytes and spermatozoa, followed by fertilization, embryo development, and implantation. Several factors influence fertility, such as the maternal age, oocyte quality, endometrial thickness, and sperm motility. Despite the utilization of assisted reproductive technology, many couples remain unable to achieve pregnancy. Research indicates that a range of genetic factors contribute to oocyte maturation arrest, fertilization failure, embryo stasis, and pre-implantation embryo demise, all of which underlie the complexity of infertility(2–5). In addition, premature ovarian insufficiency is a major cause of female infertility due to early loss of ovarian function (6, 7). It affects 3.7% of women younger than 40 (8), yet its molecular etiology is still unclear (9–11). Therefore, it is key to detect infertility early and treat childbearing-age women early to improve the quality of the population quality and fertility.

Oocyte maturation is a prerequisite for successful fertilization. Mammalian oocytes pass through the germinal vesicle (GV) stage and metaphase I (MI) stage to reach metaphase II (MII)(12, 13). In the MII stage, oocytes extrude the first polar body and gain the capacity to be fertilized(12). Engagement of mature oocytes with sperms generates zygotes, which subsequently undergo cleavage and early embryonic development before implantation(14). Oocyte maturation involves a series of nuclear and cytoplasmic maturation events(15, 16). Nuclear maturation includes germinal vesicle breaks down (GVBD), chromosome condensation and segregation, and expulsion of polar bodies. And the cytoplasmic maturation process involves the reorganization of organelles, an increase in Ca^2+^ storage contents, antioxidants(17), and the storage of mRNA and proteins(16).

As the main source of ATP and a variety of intermediates for cell metabolism (18–21), mitochondria are also involved in many other cellular processes, including oocyte maturation, defense responses to oxidative stress(22), and the regulation of programmed cell death(23). It is dynamically regulated by fusion and fission processes (24, 25), and its activity is closely related to intraorganellar interaction (26, 27). Importantly, the mitochondrial DNA (mtDNA) copy number dramatically expanded 2000-fold during oocyte maturation, increasing from approximately 200 copies per cell in primordial germ cells to 400,000 copies in matured MII oocytes(28). In addition to expansion in mtDNA copies, oocyte maturation is accompanied by changes in dynamic mitochondrial distribution and microstructure, which might be related to the increasing ATP demand, ATP production efficiency, and oxygen consumption(18, 21, 29, 30). In immature oocytes, mitochondria are clustered around germinal vesicles. As oocytes mature, mitochondria are gradually dispersed in the cytoplasm, resulting in a uniform distribution of mitochondria in the cytoplasm of MII oocytes(18, 21).

The mitochondrial dNTP pool is maintained by the dNTP salvage pathway and is critical for mtDNA replication and homeostasis, especially in terminally differentiated cells (e.g. neuronal cells) and slow cycling cells(31). Deoxyguanosine kinase (DGUOK) is a rate-limiting enzyme for the purine deoxynucleotide salvage pathway in mitochondria. DGUOK deficiency in humans leads to the hepatocerebral mtDNA deletion syndrome (MDS), which in the most severe cases, could lead to fatal infantile hepatocerebral diseases(32). Although mtDNA replication and expansion are well documented during oocyte development, the role of the mitochondrial nucleotide salvage pathway in oocyte maturation and female infertility has not been previously reported. We found that deletion of *Dguok* leads to oocyte immaturity and female infertility in mice. Our results shed new light on the mitochondrial nucleoside salvage pathway in oocyte development and female fertility.

## Materials and methods

### Animals and tissue preparation

All mice were maintained in Laboratory Animal Center of Yunnan University according to institutional guidelines. All animal procedures were approved by Institutional Animal Care and Yunnan University.

### Construction of *Dguok* knockout mouse

Establishment of *Dguok* targeted small guide RNA, consists of M-Dguok-e2be-gRNA Up, M-Dguok-E2BE-gRNA Down, M-Dguok-E2AF-gRNA Up, and M-Dguok-e2af-GrNA Down. The target is located in the intron region on both sides of the second exon of *Dguok*, nucleotide sequences include SEQ ID NO.1 and SEQ ID NO.2. sgRNA and CAS9 mRNA (CAS9MRNA-1EA, Sigma-Aldrich) were injected into the nuclear region of fertilized eggs from C57BL/6 mice with the concentration of 0.1g/L and 0.05g/L, then the embryos were transferred into the fallopian tubes of pseudopregnancy mice. The female positive founder mice were mated with wild-type C57BL/6 mice to obtain positive heterozygous F1 mice. The F1 heterozygous mice were self-crossed to obtain homozygous female F2 mice.

SEQ ID NO.1: GGCCCTGGCTCCCATGAGATGGG

SEQ ID NO.2: GGCAGGAGCAATAGTCAACGAGG

M-*Dguok*-E2BE-GrNA Up: TAGGCCCTGGCTCCCATGAGAT

M-*Dguok*-E2BE-GrNA Down: AAACATCTCATGGGAGCCAGGG

M-*Dguok*-E2AF-GrNA Up: TAGGCAGGAGCAATAGTCAACG

M-*Dguok*-E2AF-GrNA Down: AAACCGTTGACTATTGCTCCTG

### Genotype detection by PCR

Mice were numbered 21 days after birth, and cut off mouse toes, we extracted DNA using the Jacks Lab method (https://jacks-lab.mit.edu/). This DNA was used as a template for PCR amplification. The amplification primers are as follows:

M-.*Dguok*-F: 5’-CTCCCGCACTCAGTACTACAGCT-3’

M-*Dguok*-R: 5’-AGTCCAAGTCACAGGGTCCAATA-3’

M-*Dguok*-deletion: 5’-GCGACAGAACCTATAGCAGAGTG-3’

### Stimulate ovulation and in vitro fertilization

Female *Dguok* (-/-), *Dguok* (+/-), and WT mice aged 6-8 weeks were selected for ovulation induction. Pregnant Mare Serum Gonadotropin (P9970, Solarbio) was injected at 6:00 p.m at 5 IU per 10g body weight, and Human Chorionic Gonadotropin (M2530, Nanjing Aibei Biotechnology) was injected at 5 IU per 10g body weight 48h later. Female mice were killed after Human Chorionic Gonadotropin injection14h, and the ovaries and fallopian tubes were placed in petri dishes with M2 medium (M7167, Sigma). The oocytes were obtained from the ampulla of the fallopian tube and placed in the G-IVF^TM^ medium (10136, Vitrolife) for fertilization. Then, the male mouse aged 8-10 weeks with normal fertility was killed and the epididymides were placed in petri dishes with M2 medium. Puncture the epididymis to release the concentrated sperm, the sperm suspension was collected and placed in G-IVF^TM^ medium droplets (200μL) for 30 min. Sperms were collected from the edge of droplets and the concentration of sperms was calculated by Makler counting chamber. Sperm and oocytes were co-cultured for 8 hours, and the ratio of oocytes to sperm was about 1:2000. The fertilized eggs (containing 2 tripronuclears) were collected and cultured in G-1^TM^ medium (10128, Vitrolife) for 48h, and the embryos were then transferred to G-2^TM^ medium (10132, Vitrolife) for further culture to blastocysts. The culture conditions were 37°C, 6.0% CO_2_ volume concentration, and saturated humidity.

### In vitro maturation of oocytes

The oocytes in the germinal vesicle (GV) stage were collected from the follicles after Pregnant Mare Serum Gonadotropin injection 46h-48h. The oocytes in the germinal vesicle (GV) stage were transplanted into a mature culture medium for 16-18 h. The culture conditions were 37°C, 6.0% CO_2_ volume concentration, and saturated humidity. The mature culture medium was G-1^TM^ medium, and 0.1IU/ mL follicle-stimulating hormone (HOR-253, ProSpec), 0.1IU/ mL luteinizing hormone (HOR-268, ProSpec), 1μg/ mL 17β-Estradiol (E2758-250MG, Sigma-Aldrich) were added.

### Whole-mount immunofluorescence staining of oocytes

Oocytes were fixed in 4% formaldehyde at room temperature for 30 min, washed three times in PBS containing 0.1% Tween-20 (PBS-T), permeabilized for 15 min in PBS 0.2% Triton X-100 and then blocked in PBS-T with 2% BSA (Sigma) at room temperature for 1 hr, or overnight at 4°C. Primary and secondary antibodies were diluted in blocking solution, and staining was performed at room temperature for 1 hr, or overnight at 4°C. Washes after primary and secondary antibodies were done three times in PBS-T. Primary antibodies: donkey anti-mouse alpha-Tubulin 1:500 (sc-8035, Santa Cruz Biotechnologies); donkey anti-mouse Tom20 1:500 (sc-17764, Santa Cruz Biotechnologies). Secondary antibodies: donkey anti-mouse Alexa Fluor 448 (ab150105, Abcam) diluted 1:500, or donkey anti-mouse Alexa Fluor 594 (ab150108, Abcam). DAPI staining solution (AR1176, BOSTER) was added during secondary antibody incubation.

### Examination of the ROS level and the rate of natural death of Oocytes

Oocytes were collected after ovulation induction and removed granulosa cells were with hyaluronidase digestion. Then, oocytes were incubated with 5 μM Mito-SOX^TM^ Red mitochondrial superoxide indicator (Invitrogen) or DCFH-DA (beyotime) at 37°C for 30 min. The samples were washed 3 times with G-Mops^TM^ medium (10129, Vitrolife) and immediately observed under a laser scanning confocal microscope (LSM 800, Zeiss, Germany). ROS fluorescence intensity was statistically analyzed using ZEN 2.6 software. In addition, oocytes without granulosa cells were collected and placed in an M16 medium (M7292, Sigma) for further culture. The culture conditions were 37°C, 5.0% CO2 volume concentration, and saturated humidity. The rate of natural death of oocytes was counted after 6h, 30h, 54h, and 84h respectively.

### Oocyte mitochondrial electron microscope

Oocytes were fixed in 2.5% glutaraldehyde at 4℃ for 12h, then washed with PBS 3 times, and fixed at 4℃ for 1h in 1% osmic acid in the dark. Then washed with ddH_2_O. The oocytes were individually wrapped with 2% agarose gel, stained with uranyl acetate at 4℃overnight, and washed three times with ddH_2_O. Subsequently, ethanol gradient dehydration and resin gradient embedding were performed. The embedded samples were sectioned with an ultra-thin microtome, and the samples carried in the copper mesh were stained with uranyl acetate for 10min, followed by ddH_2_O washing, airtight staining with lead citrate for 5min, and then washed with ddH_2_O. After drying, multiple images of mitochondria in oocytes were taken by electron microscope, and oocyte internal structures were analyzed.

### HE staining of mouse ovaries

For histopathological analysis, the ovary was excised, washed with saline, and fixed with 4% formalin. Afterward, the tissues were embedded in paraffin and cut into 7μm-thick sections which were stained with hematoxylin and eosin (H&E). Then, sections of each mouse ovary were observed under a microscope.

### Real-Time PCR Analyses

After ovulation induction, the oocytes were collected, granulosa cells were removed with hyaluronidase solution, and single oocytes were collected into single tubes of lysis buffer (adding 1 µl of RNase inhibitor to 19 µl of a 0.2% (vol/vol) Triton X-100 solution). Reverse transcription and PCR preamplification were performed according to the previous protocol. We typically use 18 cycles for a single oocyte to obtain 10-20 ng of amplified cDNA. The RT-PCR reaction conditions were as follows: 95°C, 30 sec; 95°C, 10 sec, and 60°C, 10 sec, for 40 cycles. The system was as follows: 5μl cDNA template, 10μl 2x M5 Hiper SYBR Premix EsTaq (with Tli RNaseH) (Mei5bio), 1μl Forward primer, 1 Reverse primer, and ddH2O was supplemented to a total volume of 20μl. The fluorescence signals were analyzed by the CFX96 Real-Time PCR detection system (BIORAD). Relative expression levels of target genes were calculated with the ΔCT method using *Rpl32* or gDNA *Actb* as an endogenous reference gene for internal normalization. A total of eight *Dguok* ^-/-^ mice and four wild-type mice were ovulated and the reaction was repeated three times, total of three independent experiments.

The amplified primer pair are:

DMN1(mouse)Q-PCR Forward: 5’-AACTCTGATGCCCTCAA-3’

DMN1(mouse)Q-PCR Reverse: 5’-CTTGTTCTCTAGCACGT-3’

Miga2(mouse)Q-PCR Forward: 5’-GAGCAGGCACTGAGTGT-3’

Miga2(mouse)Q-PCR Reverse:5’-TTCTCAGCAAACTCCCT-3’

Miga1(mouse)Q-PCR Forward: 5’-ATCACAGTTCTCCTTGA-3’

Miga1(mouse)Q-PCR Reverse: 5’-ATCACAGACACAGCACT-3’

mRPL32 Q-PCR NS: 5’-ACAATGTCAAGGAGCTGGAG-3’

mRPL32 Q-PCR CAS: 5’-TTGGGATTGGTGACTCTGATG-3

QPCR_mmtDNA_Dloop1_FW:5’-AATCTACCATCCTCCGTGAAACC-3’

QPCR_mmtDNA_Dloop1_RV: 5’-TCAGTTTAGCTACCCCCAAGTTTAA-3’

gDNA Actb(M) NS: 5’-GAGACCTCAACACCCCAG-3’

gDNA Actb(M)CAS: 5’-GAGCATAGCCCTCGTAGATG-3’

### Detection of oocyte and granulosa cell apoptosis

AnnexinV-EGFP cell apoptosis detection kit (beyotime) was used to detect the level of oocyte or granulosa cell apoptosis. Oocytes obtained by stimulating mouse ovulation with hormones continued to culture for 6 hours and 36 hours, and washed with warm PBS in advance could ensure the subsequent combination of AnnexinV-EGFP. The oocytes were transferred to the pre-diluted AnnexinV-EGFP dye solution. After incubation in the dark at 37℃ for 30min, it was washed with warm 1×PBS and observed under a fluorescence microscope. Granulosa cells from the periphery of oocytes were collected and the apoptosis of granulosa cells was detected by the same procedure as that of oocytes.

### Detection of granulosa cell proliferation within the follicle

wild-type and *Dguok*^-/-^ mice were intraperitoneally injected with BrdU at a dose of 60mg/kg, and mice were sacrificed 6 hours later. Ovaries were then extracted and fixed in 10% formalin at 4°C overnight for fixation. For histology, fixed ovaries were washed, dehydrated, and embedded in paraffin. Paraffin-embedded ovaries were serially sectioned at 4 μm. The reaction was performed with mouse anti-BrdU-FITC antibody (BioLegend Cat. No. 364103) at 4℃ overnight, washed, and stained with DAPI for 20min before fluorescence observation. The extent of granulosa cell proliferation within the follicle was determined by the presence of BrdU in randomly selected ovarian sections.

## Results

### 1. *Dguok* knockout female mice are infertile

In our previous studies, we found that DGUOK deletion affected the assembly of mitochondrial complex I, and robustly inhibited lung adenocarcinoma tumor growth, metastasis, and CSC self-renewal(33, 34). We also found DGUOK regulates nicotinamide adenine dinucleotide (NAD^+^) biosynthesis independent of the mitochondrial complex I (35). To further investigate the pathophysiological function of DGUOK, we constructed *Dguok* knockout mice (Fig S1A), in which the second exon of *Dguok* was completely deleted., and the deletion of which resulted in frame-shift mutation and early termination of translation. The homozygous *Dguok* knockout mice were validated by genotyping PCR and the absence of gene expression (Fig 1A-C). *Dguok*^-/-^ mice appeared normal except for the smaller body size due to modest growth retardation (Fig 1D). The weight of *Dguok* deficient mice was about 15% lower than control (Fig 1E). They also showed a premature aging phenotype (Fig 1F), indicating that *Dguok* deficiency-mediated mitochondria dysfunction plays a key role in aging, which is consistent with previous research(36).

**Figure 1.**
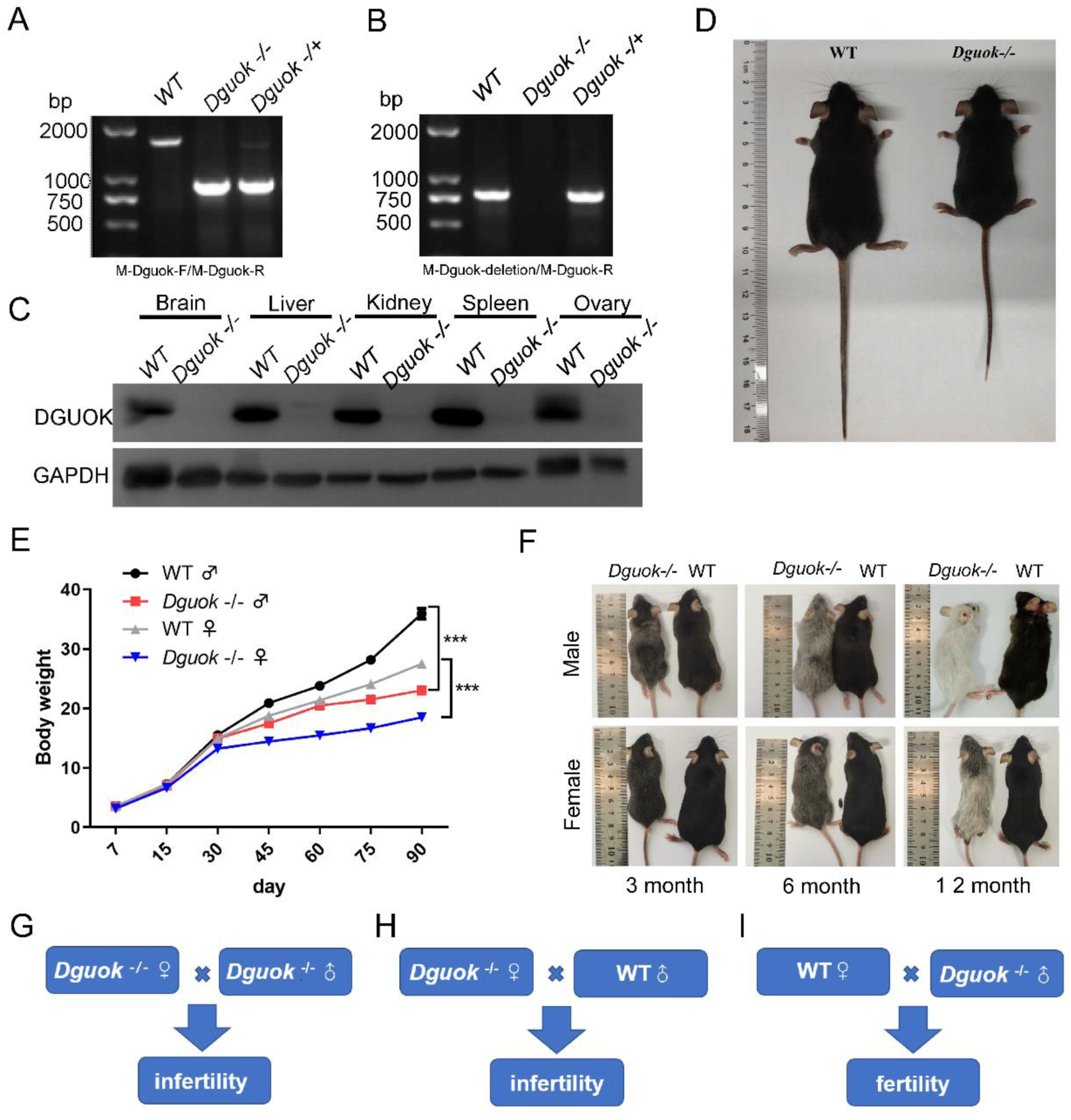
*Dguok* knockout induces female infertility. A. genotyping of *Dguok*^-/-^ mice with PCR primers M-*Dguok*-F and M-*Dguok*-R; B. genotyping of *Dguok*^-/-^ mice with PCR primers M-*Dguok*-deletion and M-*Dguok*-R; C. Western blot identification of *Dguok* expression in different tissues in WT and *Dguok*^-/-^ mice, n=3 D. Representative image of 7-weeks old female WT and *Dguok*^-/-^mice; E. Weight comparison between *Dguok-/-* and WT mice at different ages; n=10, ***p < 0.001; F. Representative image of *Dguok-/-* and WT mice at 3, 6, and 12 months. G-I. Schematic illustration of the breeding cage identifying the *Dguok*^-/-^ female’s infertility. Each breeding scheme is equipped with 10 breeding cages respectively.

Importantly, we found that mating between *Dguok*^-/-^ male mice with *Dguok^-/-^* female mice produced no progeny (Fig 1G), which suggested that DGUOK is required for fertility. We then set ten cages of mating between female *Dguok*^-/-^ mice with wild-type (WT) male mouse and found that the mating cannot produce any progeny (Fig 1H), while the mating between WT female mice with male *Dguok*^-/-^ mice produced the litter size comparable to the mating between WT C57BL/6 mice (Fig 1I). Our results suggest that homozygous *Dguok* deletion results in female infertility, but not in male mice. To investigate whether *Dguok* deficiency affects mouse gender and survival, we analyzed the offspring ratio of homozygous male mice with *Dguok* deletion to heterozygous female mice. Our results showed that the proportion of male and female and the proportion of homozygous and heterozygous in the offspring obeyed Mendel’s law. We also assessed the number of offspring of *Dguok*^-/-^ males and *Dguok*^+/-^ females. The results showed that it also obeyed Mendel’s law (Fig S1B-E). Taken together, *Dguok* knockout generates female infertility.

### 2. *Dguok* deficiency results in smaller ovaries and fewer ovulations

To investigate the causes of female infertility in *Dguok*^-/-^ mice, we examined their development of ovaries and follicles and found that their ovary size was significantly reduced compared to WT (Fig 2A, S2A). Histological analysis of the ovaries of 8-week-old female knockout mice showed numerous immature follicles, including primordial follicles with monolayer granulosa cells (Fig 2D). In comparison, the WT ovaries have more developing follicles (WT, 16.40±1.208 vs *Dguok*^-/-^,11.80±1.241) with multiple layers of granular cells (Fig 2B, 2D). Follicle development retardation was also observed in the ovaries of *Dguok*^-/-^ mice after ovulation induced by pregnant mare’s serum gonadotrophin (PMSG) and human chorionic gonadotrophin (HCG), where most oocytes were not expelled, indicating impaired response to hormones. In comparison, WT ovaries had more corpus hemorrhagicum (WT, 6.20±0.37 vs *Dguok*^-/-^, 2.33±0. 67) and more excreted oocytes (WT, 26.82±1.80vs *Dguok*^-/-^,17.27±1.46) (Fig 2C, 2E, and S2C). In sum, the ovaries from *Dguok*^-/-^ mice were hypoplastic, and the number of developing follicles and ovulations was significantly reduced (Fig 2D, S2B).

**Figure 2.**
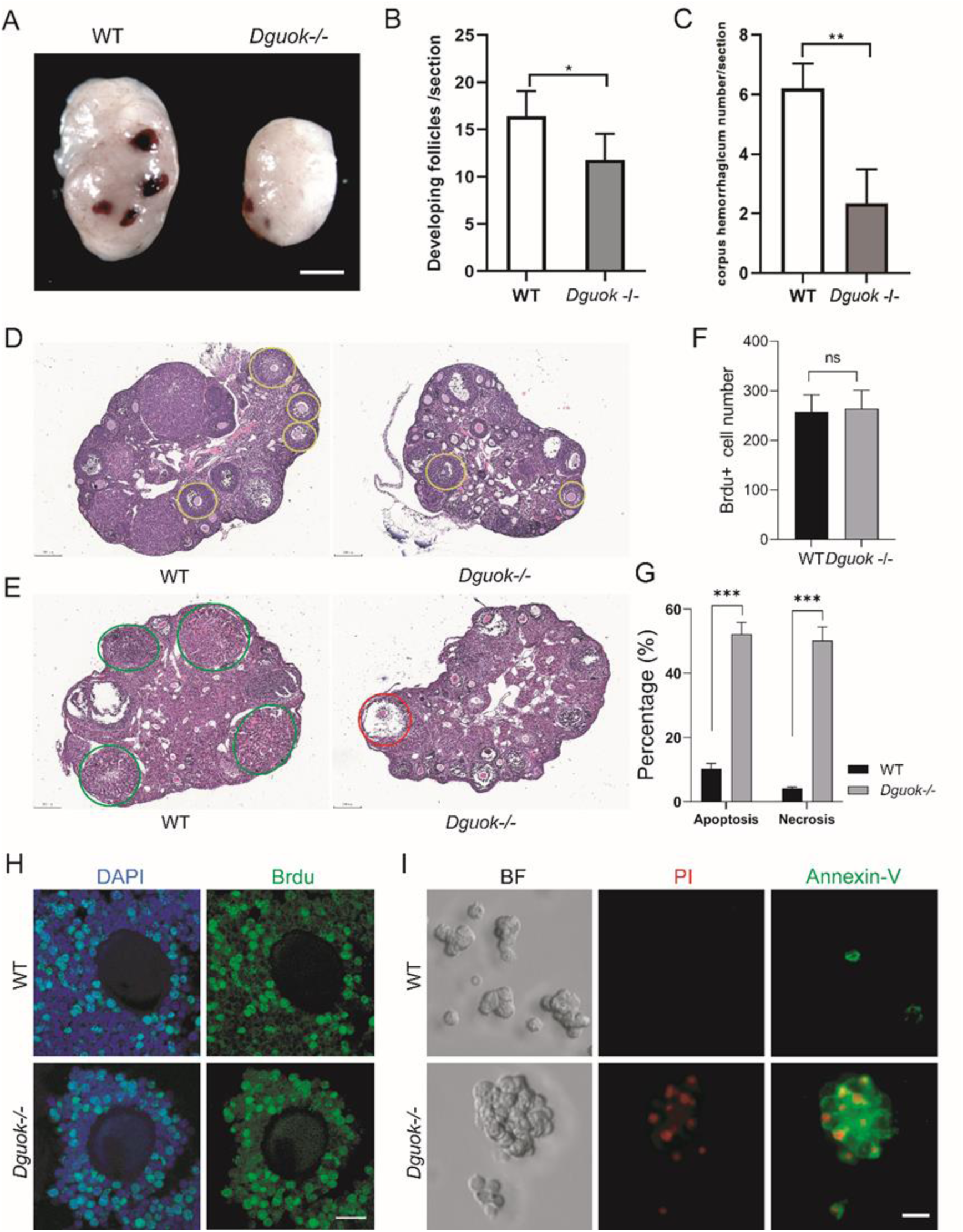
DGUOK deficiency results in smaller ovaries and fewer ovulations. A. Representative image of ovary morphology between WT and *Dguok*-/- mice, Scale bar, 1 mm; B-C. The number of developing follicles (B) and corpus hemorrhaglcum (C) per section was counted in WT and *Dguok*-/- mice; n = 3, *P<0.05 and **P< 0.01. D. Representative image of HE staining of ovarian structure, ovaries from 6-weeks-old WT and *Dguok*-/- mice were embedded, sectioned, and stained with hematoxylin and eosin (H&E), developing follicles were indicated by yellow circle, Scale bars,200μm; E. Representative image of H&E staining of ovarian structures in 6-weeks-old WT and *Dguok*-/- mice stimulated by PMSG and HCG, corpus hemorrhagicum (green circle) from WT and anovulatory dominant follicles from *Dguok*-/- mice (red circle) are indicated, Scale bar, 200 μm; F. The BrdU+ cell number of developing follicles was counted; n = 3, ns (not significant) indicates P > 0.05. G. Statistical analysis of the number of apoptotic and necrotic granulosa cells in WT and *Dguok*-/- mice; n = 3, *P<0.05 and **P< 0.01. H. Representative immunofluorescence of WT and *Dguok*-/- ovarian sections stained with BrdU antibodies; n = 3. Scale bars, 20 μm; I. Representative images of Annexin-V/PI staining indicating granulosa cell apoptosis and necrosis in WT and *Dguok*-/- mice. n = 3. Scale bars, 20 μm;

During follicular development, oocyte maturation mainly depends on the regulation of energy substrates and multiple signals provided by granulosa cells (37, 38). Therefore, the effects of *Dguok* deletion on granulosa cell proliferation and apoptosis were also examined. We found that granulosa cell proliferation (WT, 258±19.50 vs *Dguok*^-/-^,264±21.55) within the primary or secondary follicles in *Dguok^-/-^* mice was not significantly different from WT (Fig 2F, 2H, and S2D-E). However, the percentage of granulosa cell apoptosis in *Dguok^-/-^* mice was significantly higher than that in WT (Fig 2G, 2I, and S2F), which explains the significant reduction of granulosa cells around oocytes in *Dguok^-/-^* mice. Taken together, our results indicate DGUOK deficiency caused abnormal development of ovaries and oocytes.

To exclude the possibility that the infertility was due to an abnormal internal fertilization environment, we performed *in vitro* fertilization. First, we found that the size of the cumulus-oocyte complexes from *Dguok*^-/-^ mice was significantly smaller than WT. The corona was loose, and the granulosa cells in the periphery were detached (Fig S2B). Moreover, the *Dguok*^-/-^ oocytes failed *in vitro* fertilization (Fig 3A, S3A). Interestingly, we found that the majority of *Dguok*^-/-^ oocytes were arrested at the germinal vesicle (GV) stage even after 8 h of co-incubation of sperms and oocytes (Fig 3A). Meanwhile, there were no significant differences in the oocyte maturation rate, fertilization rate, and embryonic development potential between heterozygous and WT mice (Fig S3B). Together we inferred that the infertility of the *Dguok*^-/-^ mice was due to oocyte immaturity.

**Figure 3.**
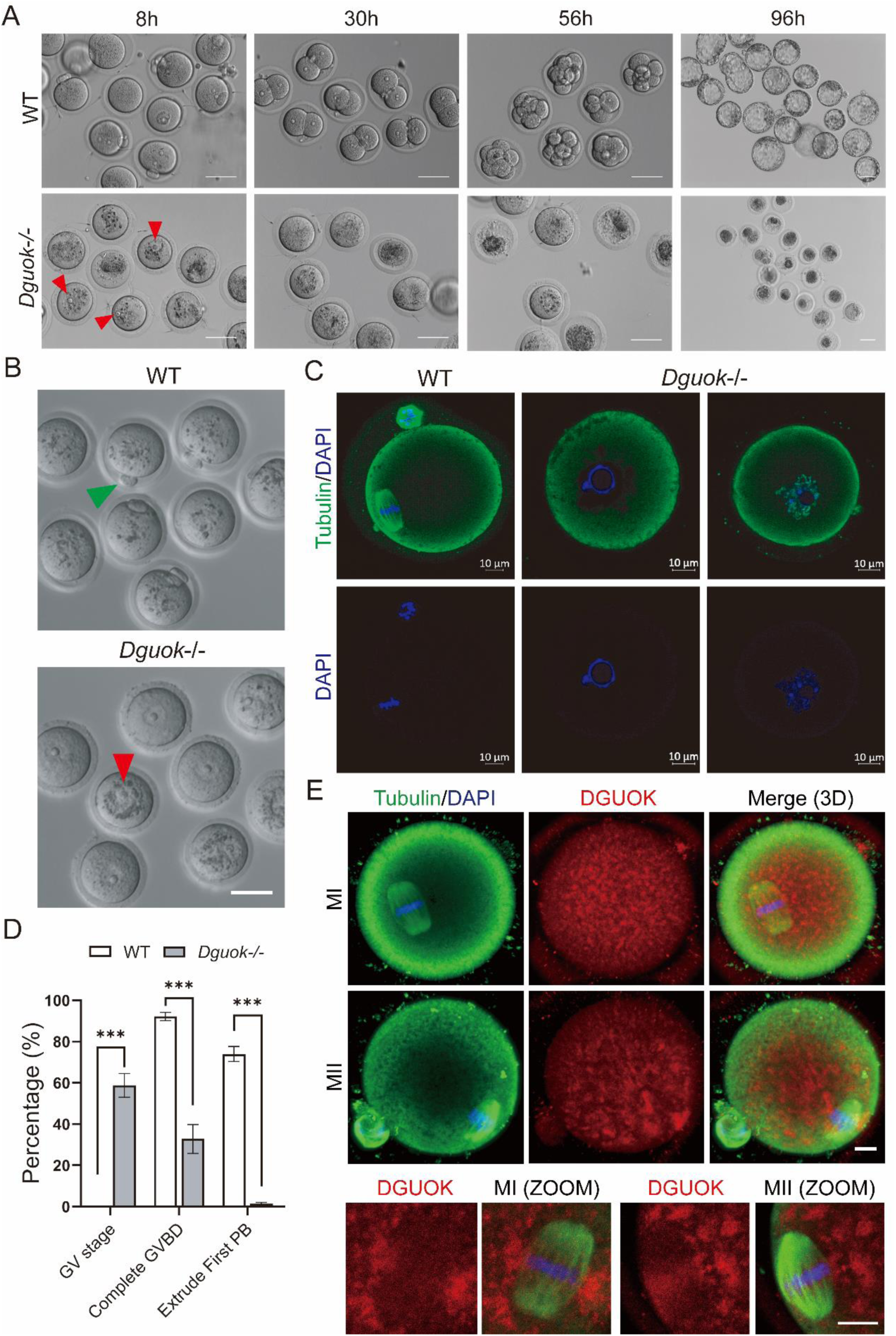
DGUOK regulates mouse oocyte meiotic maturation. A. Representative image of fertilization, 2-cells, 8-cells, and blastocyst formation after in vitro fertilization from WT and *Dguok*-/- mice; red arrows indicate GV oocytes; n > 3. Scale bars, 50 μm. B. Representative images of oocytes from WT and *Dguok*^-/-^mice. The green arrow indicates that the first polar body oocyte is expelled, red arrows indicate GV oocytes; n > 3. Scale bars, 50 μm. C. Representative images of tubulin staining in MII oocytes from WT mouse, in GV or Pre-MI oocytes from *Dguok*^-/-^ mouse. Immunofluorescence staining was performed with antibodies against tubulin (green) and DAPI (blue); n=3, Scale bars: 10 μm; D. Quantification of maturation rate *in vivo* of oocytes from WT and *Dguok*^-/-^ mice; n = 3, ***P < 0.001. E. Representative images of oocytes from WT mice for DGUOK and tubulin staining. Immunofluorescence staining was performed with antibodies against tubulin (green), DGUOK (red), and DAPI (blue); n=3, Scale bars: 10 μm.

### 3. DGUOK regulates oocyte meiotic maturation

To further investigate the effect of DGUOK on oocyte meiotic maturation, we removed the granulosa cells surrounding the *Dguok*^-/-^ oocyte and examined the spindles and chromosomes to determine their meiotic progression. Interestingly, approximately 60% of the *Dguok*^-/-^ oocytes were blocked at the GV stage and failed to resume meiosis (Fig 3B-D), while about 37% of the *Dguok*^-/-^ oocytes completed GVBD and were blocked at prophase I (Fig 3B-D). Only about 3% of *Dguok*^-/-^ oocytes extruded the first polar body (Fig 3E). In comparison, approximately 70% of WT oocytes excluded the first polar body and reached metaphase II, and no oocytes were arrested at the GV stage (Fig 3B-D). Meanwhile, we examined the distribution of DGUOK in WT oocytes. Immunofluorescence results showed that DGUOK was uniformly distributed in the cytoplasm of MI and MII oocytes, but it was more densely distributed around the spindle (Fig 3E).

Furthermore, *in vitro* oocyte maturation culture was performed to investigate the effect of intra-follicular environment on oocyte maturation. we collected oocytes at the GV stage and *in vitro* cultured them as previously described(39). We found that after 16 h of *in vitro* maturation culturing, approximately 51% of *Dguok^-/-^* oocytes completed GVBD (Fig 4B), which was significantly lower than WT (∼91%). Meanwhile, approximately 40% of *Dguok^-/-^* oocytes remained arrested in the GV stage (Fig 4B), which was significantly higher than WT (∼7.8%). Fortuitously, about 5.7% of the *Dguok*^-/-^ oocytes extruded the first pole body (Fig 4A-B, S4A), which was significantly lower than WT (∼63%). Importantly, even among the *Dguok*^-/-^ oocytes with the first pole body, the structure of the spindle apparatus was irregular, and the chromosomes were not properly aligned (Fig 4C-D, S4B), suggesting that the *in vitro* culture environment promotes the recovery of meiosis in *Dguok*^-/-^ oocytes to some extent. Our results revealed that DGUOK regulates mouse oocyte meiotic maturation both *in vivo* and *in vitro*.

**Figure 4.**
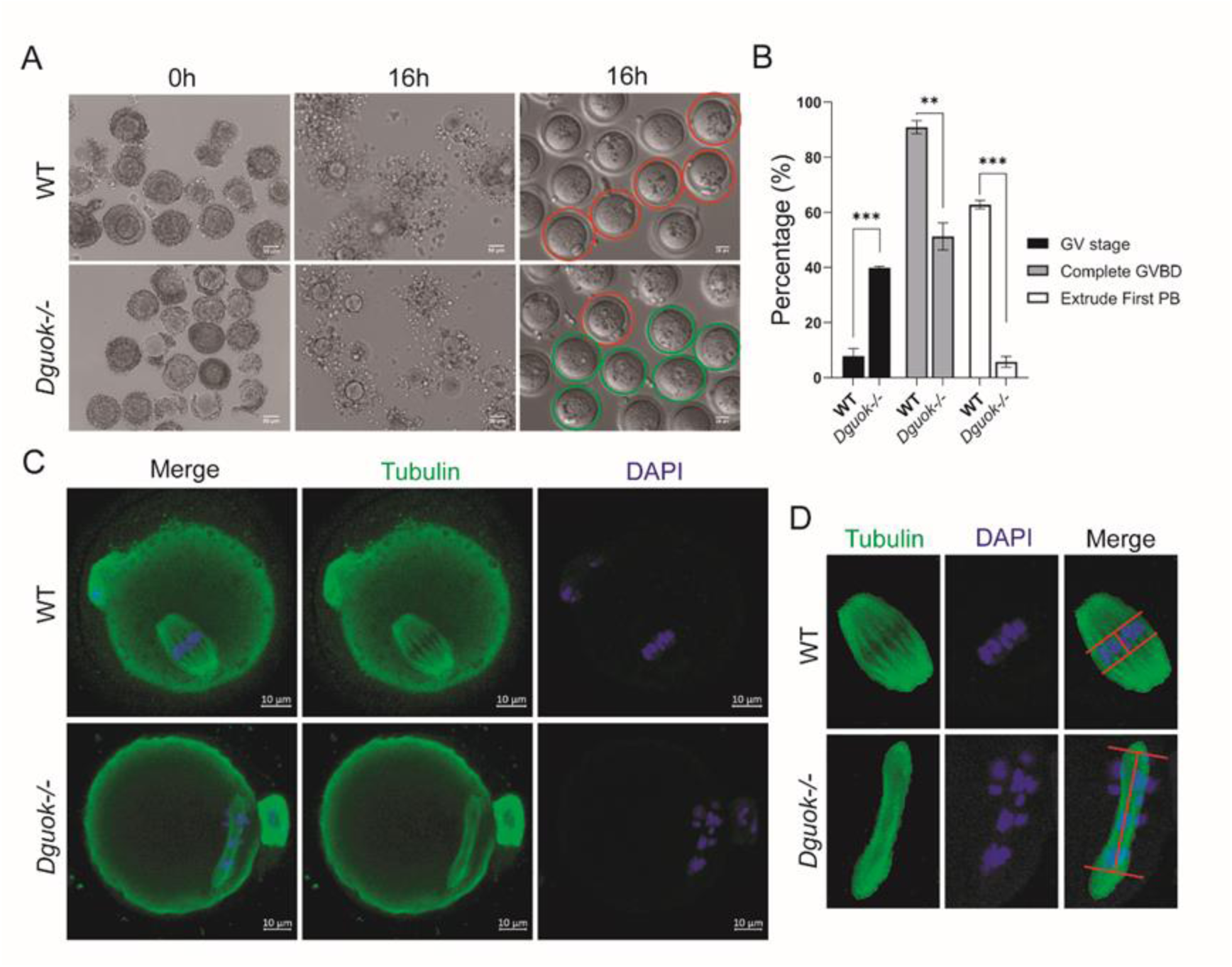
DGUOK deficiency results in irregular spindle structure and disordered chromosome arrangement. A. Representative images of cumulus-oocyte complexes mature *in vitro* (0 h, 16 h, naked-oocyte) from WT and *Dguok*^-/-^ mice; the first polar body oocyte (red circle) and GV oocytes (green circle) are indicated, n=3, Scale bars: 20 μm; B. Quantification of the GV ratio, GVBD ratio, and first pole body exclusion ratio from WT and *Dguok*^-/-^ mice after 16 h in vitro maturation; n = 3. **P<0.005 and ***P < 0.001. C. Representative images of tubulin staining in MII oocytes from WT and *Dguok*^-/-^ mice after 16 h in vitro maturation. Immunofluorescence staining was performed with antibodies against tubulin (green) and DAPI (blue); Scale bars: 10 μm; D. Enlarged images of oocyte spindle bodies from WT and *Dguok*^-/-^ mice after 16 h in vitro maturation. The red lines represent chromosome arrangement intervals. Scale bars: 10 μm;

### 4. DGUOK is essential for mitochondrial functions in mouse oocytes

Mitochondrion harbors its mtDNA, which encodes 13 critical proteins for the assembly and activity of mitochondrial respiratory complexes(40). The mtDNA plays important roles during oogenesis and early embryo development(41). Therefore, we analyzed the mtDNA copy number in *Dguok^-/-^*and WT oocytes. The results showed that WT oocytes had a significant increase in mtDNA copy number from the GV stage to the MII stage (Fig 5A). However, in *Dguok^-/-^* oocytes, the mtDNA copy number was significantly lower than that in WT oocytes at both GV and GVBD stages (Fig 5B-C). Furthermore, we also measured the mtDNA copy number in different tissues of WT and *Dguok-/-* mice and found that in addition to the significantly reduced mtDNA copy number in the liver, brain, kidney, and spleen of *Dguok^-/-^* mice, the mtDNA content in their ovaries was also severely reduced (Fig S5A). These results suggest that the deletion of DGUOK severely affects mtDNA replication, which in turn affects mitochondrial function.

**Figure 5.**
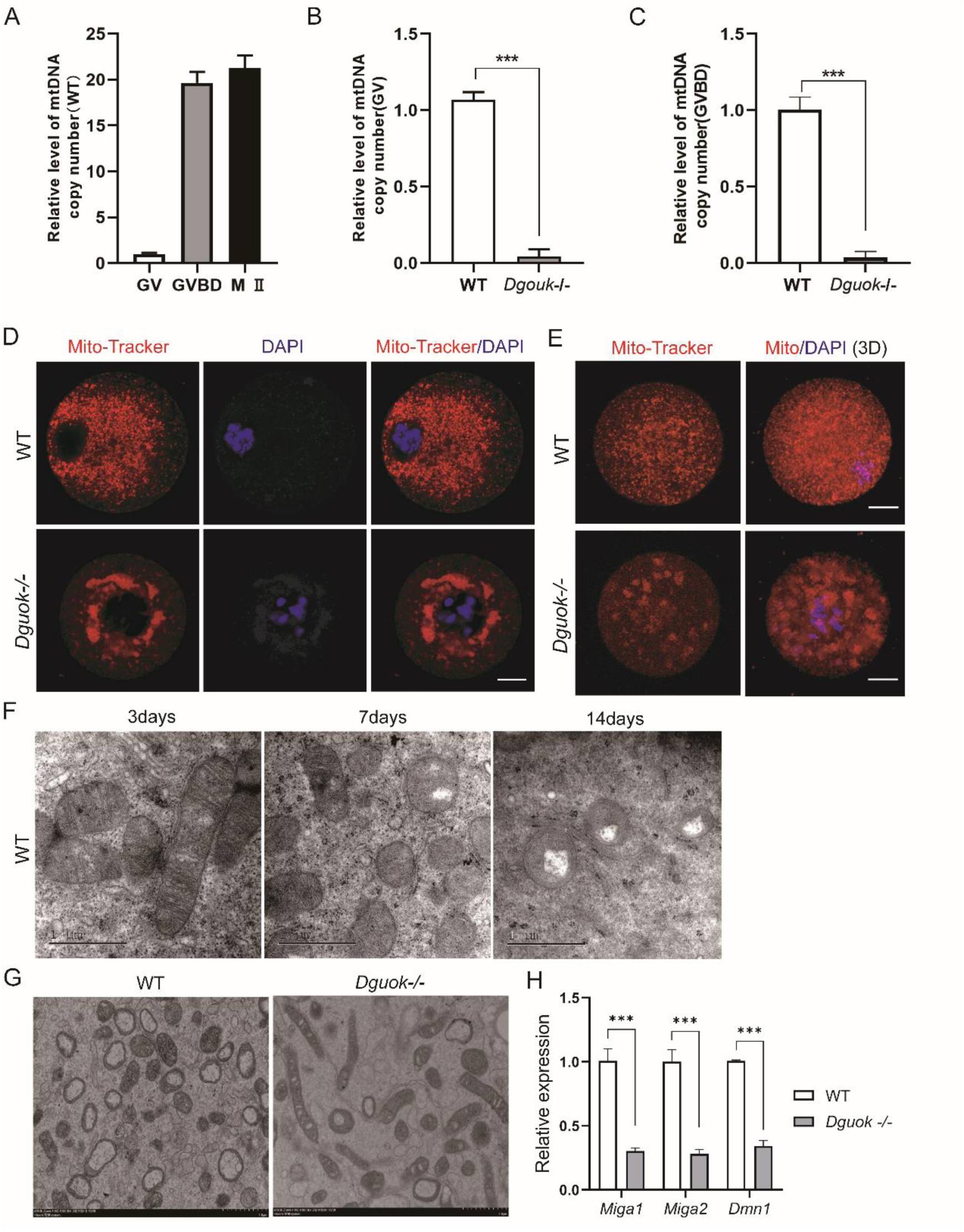
DGUOK is essential for mitochondrial functions in mouse oocytes. A. mtDNA content single oocytes of WT mouse at different stages, n=3; B. Relative level of mtDNA copy number in GV oocytes from WT and *Dguok*^-/-^ mice, n=3, ***p < 0.001. C. Relative level of mtDNA copy number in GVBD oocytes from WT and *Dguok*^-/-^ mice, n=3, ***p < 0.001. D. Representative images of Mito-Tracker staining in WT and *Dguok*^-/-^ GVBD oocytes; Scale bars: 20 μm. E. Mito-Tracker staining in the oocytes from WT and *Dguok*-/- mice with PMSG and HCG stimulation; Scale bars: 20 μm. F. TEM images of oocyte mitochondria in the ovaries of WT mice at 3, 7, and 14 days after birth; Scale bars: 1 μm. G. Representative mitochondria TEM images of oocytes from WT and *Dguok*^-/-^ mice; Scale bars: 1 μm. H. mRNA expression levels of *Miga1, Miga2, Dmn1* in oocytes of WT and *Dguok*^-/-^ mice, n=3, ***p< 0.001.

The distribution of mitochondria in oocytes changes dynamically with the energy demand of oocytes at different stages of development. At the GV stage, mitochondria in the oocyte gradually accumulate in the perinuclear space to provide energy for nuclear rupture and spindle traction. Subsequently, mitochondria gradually distribute evenly throughout the oocytes to maintain the basal metabolism level (21, 42, 43). However, in *Dguok^-/-^* oocytes, the movement and distribution of mitochondria are abnormal: mitochondria were irregularly clustered around the nucleus and in the cytoplasm, both before and after the germinal vesicle breakdown (Fig 5D-E). Meanwhile, we also analyzed the ultrastructure of *Dguok*^-/-^ and WT oocytes by electron microscopy. We found that mitochondria gathered in clusters in *Dguok*^-/-^ oocytes (Fig S5C), which was consistent with the immunofluorescence staining results (Fig 5D-E). Therefore, the deletion of DGUOK leads to an imbalance in the dynamic distribution of mitochondria in oocytes, which interferes with the function of mitochondria during oocyte maturation.

It is known that mitochondria change their position and morphological structures (such as elongation, shortening, and swelling) in cells through fusion and fission to maintain mitochondrial homeostasis and normal function(44, 45), which are essential for providing energy, countering stress, repairing mtDNA and mitosis (46–48). It has been reported that mitochondria in the oocytes of neonatal mice are mainly elongated dumbbells, but as the mice grow, mitochondria in the oocytes become round and oval (49). To further investigate whether deletion of *DGUOK* causes abnormal changes in the mitochondria dynamics in oocytes, we analyzed the morphology of mitochondria in oocytes of *Dguok*^-/-^ and WT mouse at 3, 7, or 14 days after birth by electron microscopy. Our results showed that mitochondria were elongated dumbbells with transverse cristoids from 3-day-old WT mouse oocytes (Fig 5F, S5D), but it became round and vacuolar at days 7 and 14 of birth (Fig 5F, S5D) and until they reached adulthood (8-week-old) (Fig 5G, S5E). In contrast, the mitochondrial morphology of *Dguok*^-/-^ oocytes (from 8-week-old mice) showed elongated dumbbells (Fig5G, S5E). Meanwhile, we examined the expression of genes critical for mitochondrial fission regulation in oocytes (50, 51). The expression of *Miga1*, *Miga2*, and *Dmn1* in *Dguok*-/- oocytes was significantly decreased than WT (Fig 5H). Therefore, deletion of *DGUOK* impaired mitochondrial fission function in oocytes. Taken together, we conclude that DGUOK is essential for mitochondrial homeostasis and function during oocyte maturation.

### 5. DGUOK deficiency significantly increased reactive oxygen species (ROS) and apoptosis of oocytes

Mitochondria are the metabolic hubs and the main source of ROS. High levels of ROS cause deleterious damage to the cell and gene structure and may lead to apoptosis. We found that *Dguok*^-/-^ oocytes had a higher necrosis rate than WT in *in vitro* culture medium (Fig S6A-B), indicating increased oocyte apoptosis in *Dguok*^-/-^ oocytes. Subsequently, we used the Annexin-V apoptosis detection reagent to stain the oocytes maintained *in vitro* for 6 h and 36 h (Fig 6A). As expected, the apoptosis rate of *Dguok*^-/-^ oocytes was significantly higher than WT (Fig 6A). Meanwhile, we detected ROS levels using DCFH-DA and Mito-sox staining in oocytes. Fluorescence intensity analysis showed that the content of mitochondrial ROS in *Dguok*^-/-^ oocytes was significantly higher than WT (Fig 6B-D). This revealed that DGUOK deficiency led to significantly higher ROS levels and increased apoptosis in oocytes.

**Figure 6.**
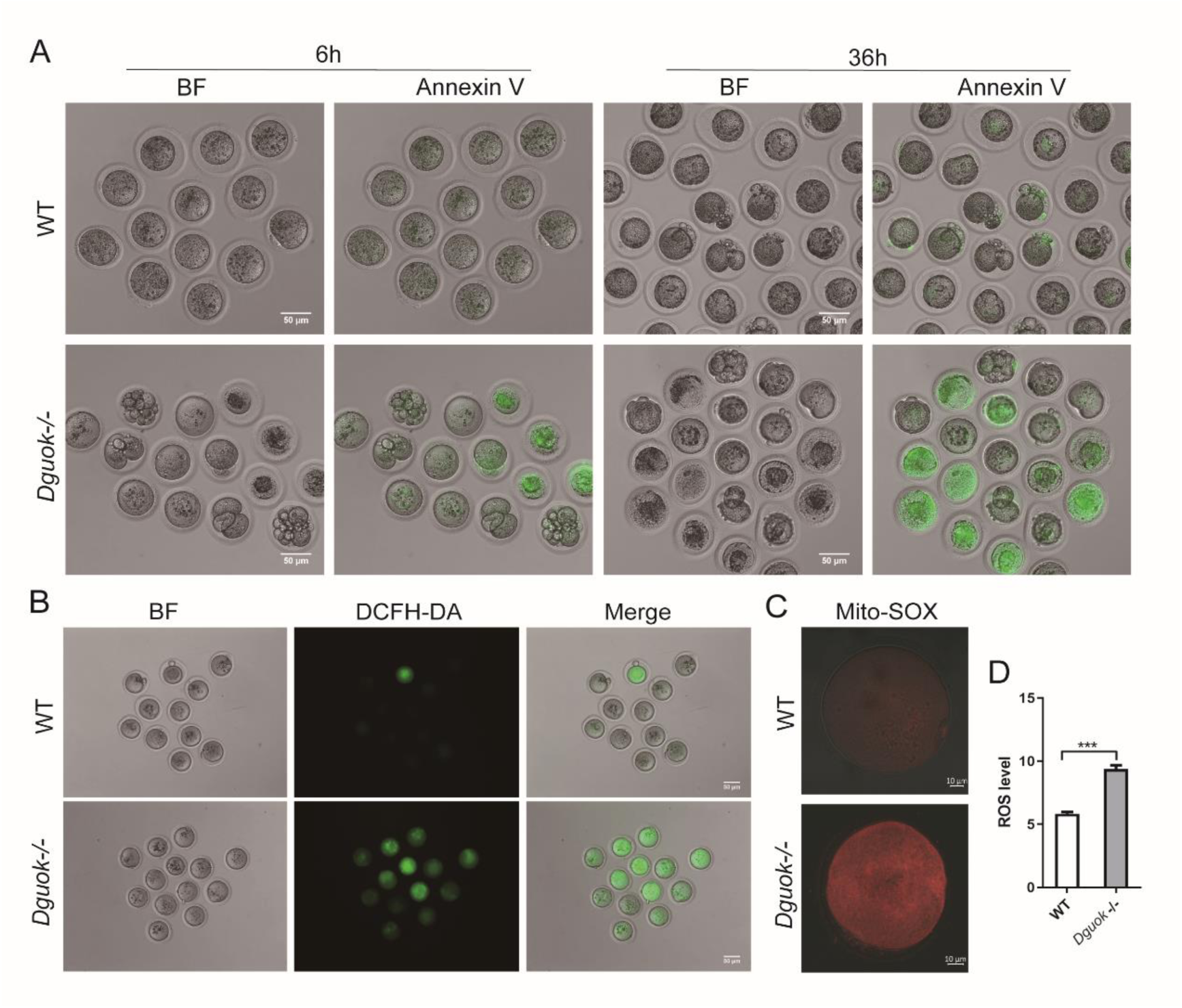
DGUOK deficiency significantly increased ROS and apoptosis of oocytes. A. Annexin-V staining in oocytes (maintained in vitro for 6 h and 36 h) from WT and *Dguok*^-/-^ mouse; Scale bars: 50 μm; B. Representative image of DCFH-DA signals in oocytes from WT and *Dguok*^-/-^ mouse; Scale bars: 50 μm; C-D. Representative images and quantification analysis of ROS fluorescence intensity in oocytes from WT and *Dguok*^-/-^ mouse, n>3, ***p < 0.001. Scale bars: 10 μm in C.

## Discussion

In this study, we found that DGUOK deficiency seriously disrupts mitochondrial homeostasis and function in oocytes. The dysfunction of mitochondria causes an increase in ROS levels, apoptosis of oocytes, and seriously disturbs the meiotic process of oocytes, ultimately leads to oocyte immaturity. Interestingly, we found that *Dguok* deletion induces female mice infertility but not in males. Moreover, DGUOK*-*mediated mitochondria dysfunction plays important roles in oocyte maturation and Infertility.

Oocyte maturation requires many energy-demanding processes such as cytoskeleton remodeling, chromosome synapsis, vesicle transport, and membrane resynthesis (52). Mitochondria are the key regulators of multiple vital cellular processes including oocyte maturation, and mitochondria provide the energy source and signal transduction for oocyte development. Mitochondrial dysfunction will fail to provide energy and other support for the proceeding of these cellular processes (53). In *Dguok*^-/-^ mice, oocytes failed to complete meiosis, and most of them were arrested at the GV stage, indicating that mitochondria play an important role in oocyte meiosis. However, we cannot exclude the possibility that the expression of some key mitochondrial genes is impaired in *Dguok*^-/-^ mice, which may cause meiotic arrest in oocytes.

Cytoplasmic maturation of oocytes is indispensable for fertilization. Mitochondria are inherited from the maternal cytoplasm and are potential contributors to cytoplasmic maturation. The mtDNA copy number was reported to correlate with the potential of the human oocytes to be fertilized(54). It has been reported that infertility at an advanced maternal age is associated with mtDNA mutation, suggesting that mtDNA plays key roles in infertility(28), and may serve as a potential biomarker for embryo viability in assisted reproduction(55, 56). Thus, the correlation between mitochondrial and reproduction, particularly in oocyte maturation, is underlined, though the underlying mechanism is unclear due to lack of optimal tools.

In the process of follicular development, the development and maturation of oocytes mainly depend on the provision of energy substrates by granulosa cells(37, 38). Meanwhile, in the process of oocyte maturation, granulosa cells transmit a variety of signals through gap junctions, which participate in the initiation of oocyte meiosis and the maturation of oocyte cytoplasm(57). The autocrine and paracrine substances of granulosa cells can promote the proliferation of granulosa cells and the growth of follicles. At the same time, the interaction between granulosa cells and thecal cells is important for the development and maintenance of normal functions of follicles. Therefore, granulosa cells play an important regulatory role in the initiation, growth, and atresia of primordial follicles (58). We found that the number of growing follicles in the ovaries of *Dguok*-/- mice was significantly less than WT, while the number of atretic follicles was significantly increased, and the number of granulosa cells in the periphery of oocytes was significantly less than WT, which may be related to the abnormal growth, differentiation, and apoptosis of granulosa cells in *Dguok*^-/-^ mice. Due to the loss of DGUOK, the abnormal mitochondrial function of granulosa cells may lead to the abnormal follicular microenvironment, which in turn inhibits the development and maturation of oocytes. These may be important reasons why *Dguok^-/-^* oocytes are arrested at the GV stage *in vivo* but can partially resume meiosis in mature culture *in vitro*. In addition, the ovarian insufficiency caused by *Dguok* deficiency also provides a potential molecular target for the pathogenesis of primary ovarian insufficiency (POI).

Taken together, our data support that DGUOK regulates oocyte maturation and maternal reproduction. Future investigations to identify whether *Dguok* affects maternal factors are required. Our results thus provide therapeutic strategies for the treatment of infertility and a target for prenatal diagnosis.

## Acknowledgment

We thank Dr. Jian Zhang for the helpful suggestion, reagents, and comments on the manuscript. We also thank Dr. Jing Li for the critical reading of the manuscript. We also thank the Animal Research and Resource Center of Yunnan University for the mice breeding management and experimental operation service.

---

**Figure S1.**
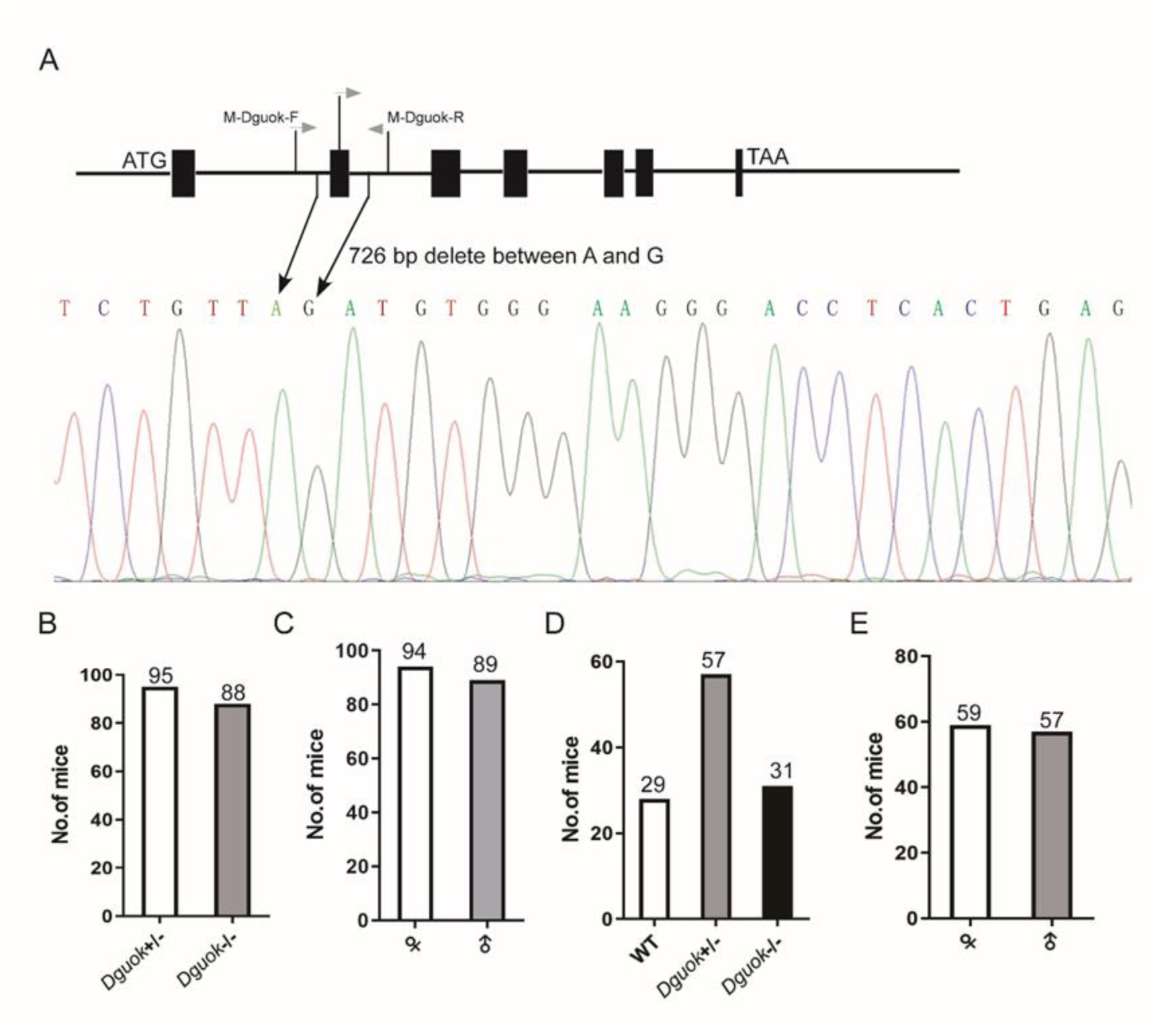
Construction of Dguok knockout mouse. A. scheme of 726 bp deletion in *Dguok* knockout mice; B. statistical number of offspring of *Dguok*^-/-^ male with *Dguok*^+/-^ female; C. Statistical number of male and female offspring of *Dguok*^-/-^ male with *Dguok*^+/-^ female; D. Statistical the number of wild: heterozygous: homozygous offspring of *Dguok*^+/-^ inbred; E. The statistical number of male and female offspring of *Dguok*^+/-^ inbred; each breeding scheme is equipped with 10 breeding cages respectively.

**Figure S2.**
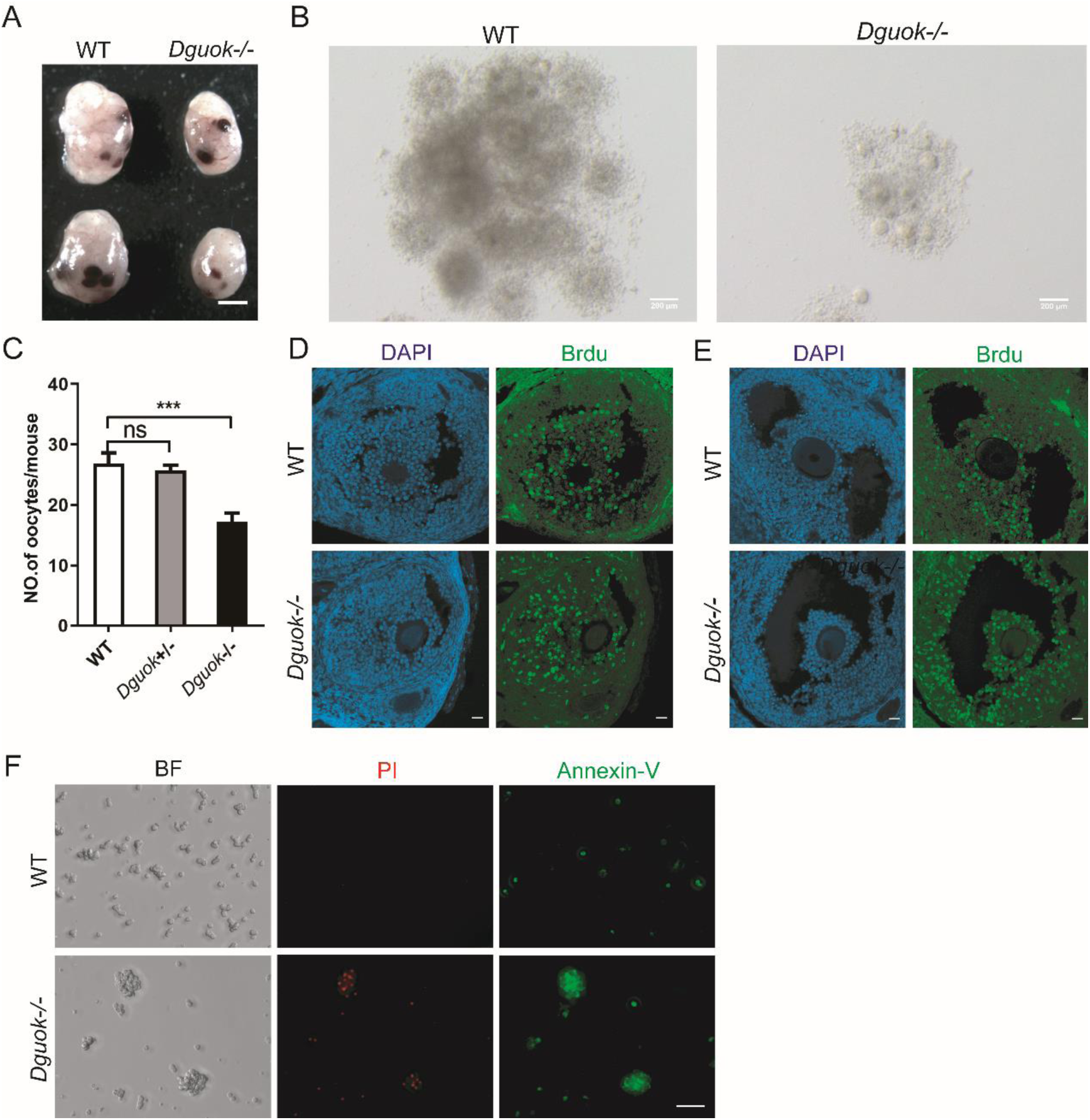
DGUOK deficiency results in smaller ovaries and fewer ovulations. A. Representative image of ovary morphology between WT and *Dguok*^-/-^ mice; Scale bars: 1mm in A; B. Representative image of cumulus-oocyte complexes in WT and *Dguok*^-/-^ mice; Scale bars: 200μm; C. Statistics on the number of oocytes from WT and Dguok^-/-^ mice; n > 3, ns indicates P>0.05, ***P < 0.001. D. Representative immunofluorescence micrographs of primary follicles from WT and *Dguok*^-/-^ ovarian sections stained with BrdU-FITC antibody (green) and DAPI (blue); Scale bars: 20μm; E. Representative immunofluorescence micrographs of secondary follicles from WT and *Dguok*^-/-^ ovarian sections stained with BrdU-FITC antibody (green) and DAPI (blue); Scale bars: 20μm; F. Representative images of Annexin-V/PI staining indicating granulosa cell apoptosis and necrosis in WT and *Dguok*^-/-^ mice; Scale bars: 50μm;

**Figure S3.**
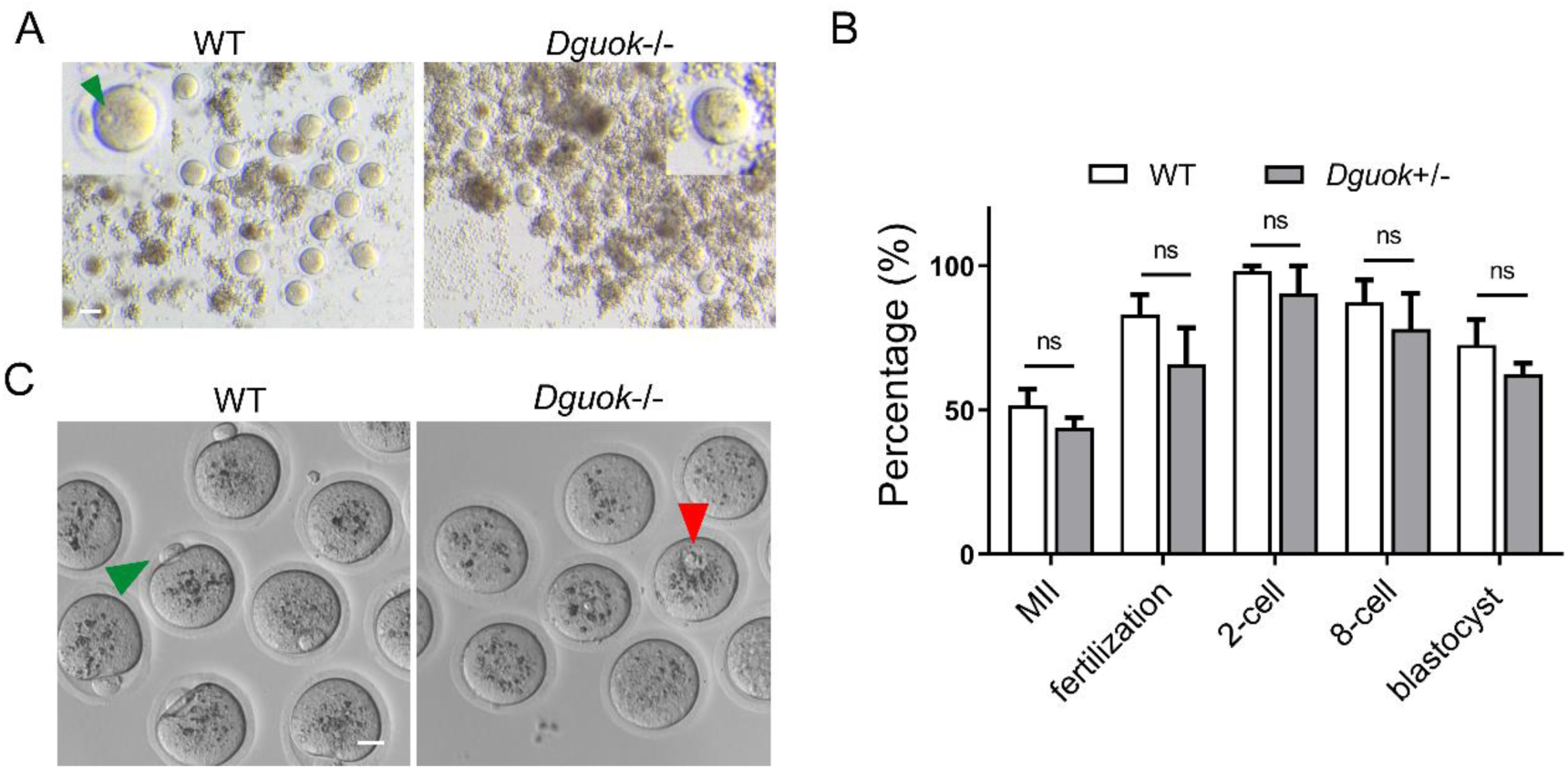
DGUOK is essential for mouse oocyte meiotic maturation. A. Representative images from WT and *Dguok*^-/-^ mice oocytes after IVF, green arrows indicate male pronucleus, n>3,Scale bars: 50μm. B. Quantification of stages in MII, fertilization, 2-cells, 8-cells, and blastocyst from WT and *Dguok*^+/-^ mice oocyte after IVF, n>3, ns indicates p > 0.05. C. Representative image of oocytes after IVF from WT and *Dguok*^-/-^ mice, The green arrow indicates that the first polar body oocyte is expelled, and red arrows indicate GV oocytes. n>3; Scale bars: 20μm.

**Figure S4.**
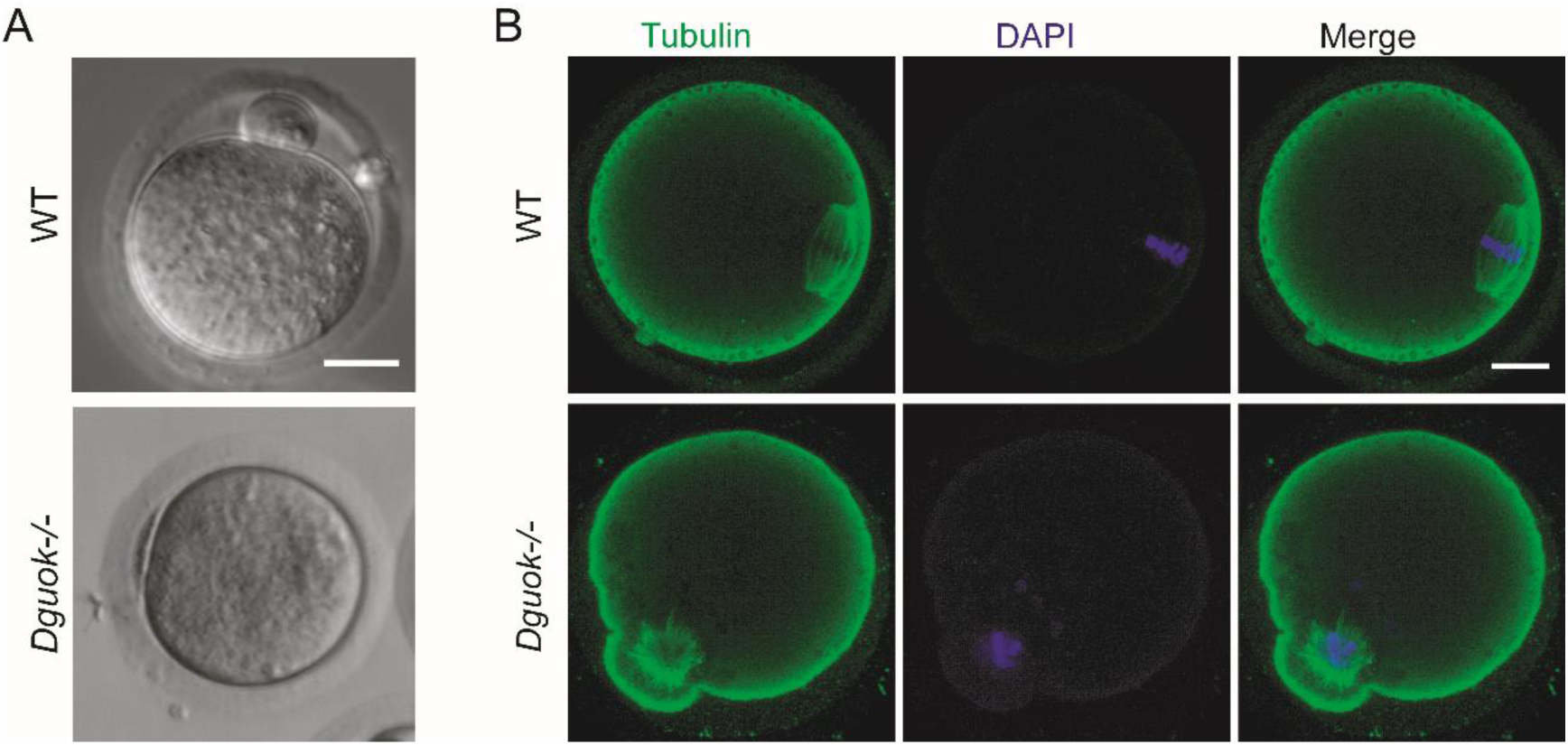
DGUOK deficiency results in abnormal meiosis of oocytes. A. Representative images from WT and *Dguok*^-/-^ mice oocytes after mature in vitro; Scale bars: 20μm B. Representative image of tubulin staining in MII-like oocyte from WT and *Dguok*^-/-^ mice after 16 h in vitro mature. Immunofluorescence with antibodies against Tubulin (green) and DAPI (blue); Scale bars: 10μm.

**Figure S5.**
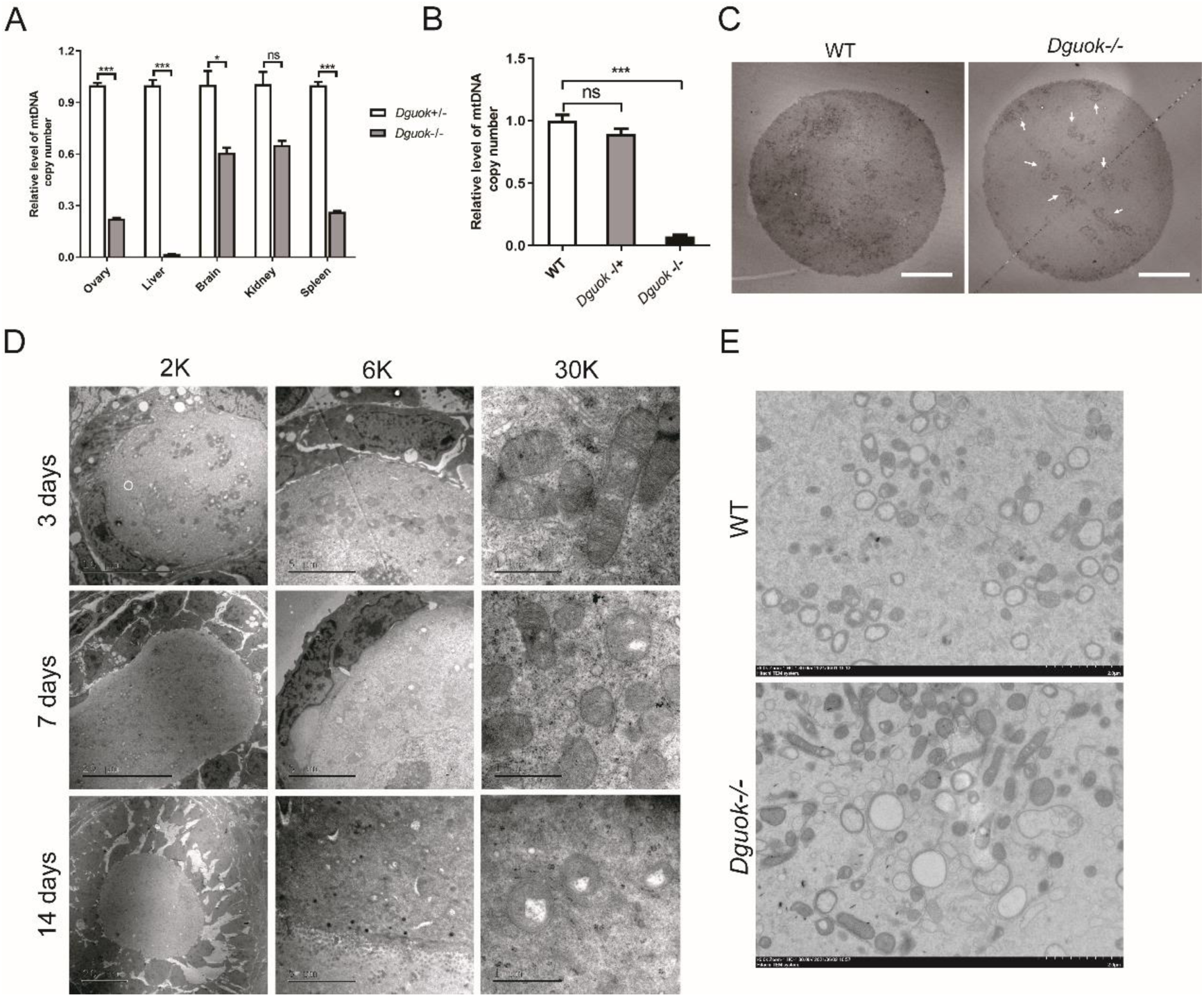
DGUOK is essential for mitochondrial functions in mouse oocytes. A. Relative level of mtDNA copy number in different tissues of WT and *Dguok*^-/-^ mice; n=3; ns indicates p > 0.05, *p < 0.05, ***p < 0.001. B. Relative level of mtDNA copy number in oocytes from WT, *Dguok^-/+^,* and *Dguok*^-/-^ mice; n=3; ns indicates p > 0.05, *p < 0.05, ***p < 0.001. C. Representative Mitochondrial distribution TEM images of oocytes from WT and *Dguok*^-/-^ mice, white arrows indicate mitochondrial aggregation Scale bars: 20μm; D. TEM images of oocyte mitochondria in the ovaries of WT mice at 3 days, 7 days, and 14 days after birth, the magnification was 2k, 6k, 30k; Scale bars: 1μm; E. Representative mitochondria TEM images of oocytes from WT and *Dguok*^-/-^ mice.Scale bars: 1μm;

**Figure S6.**
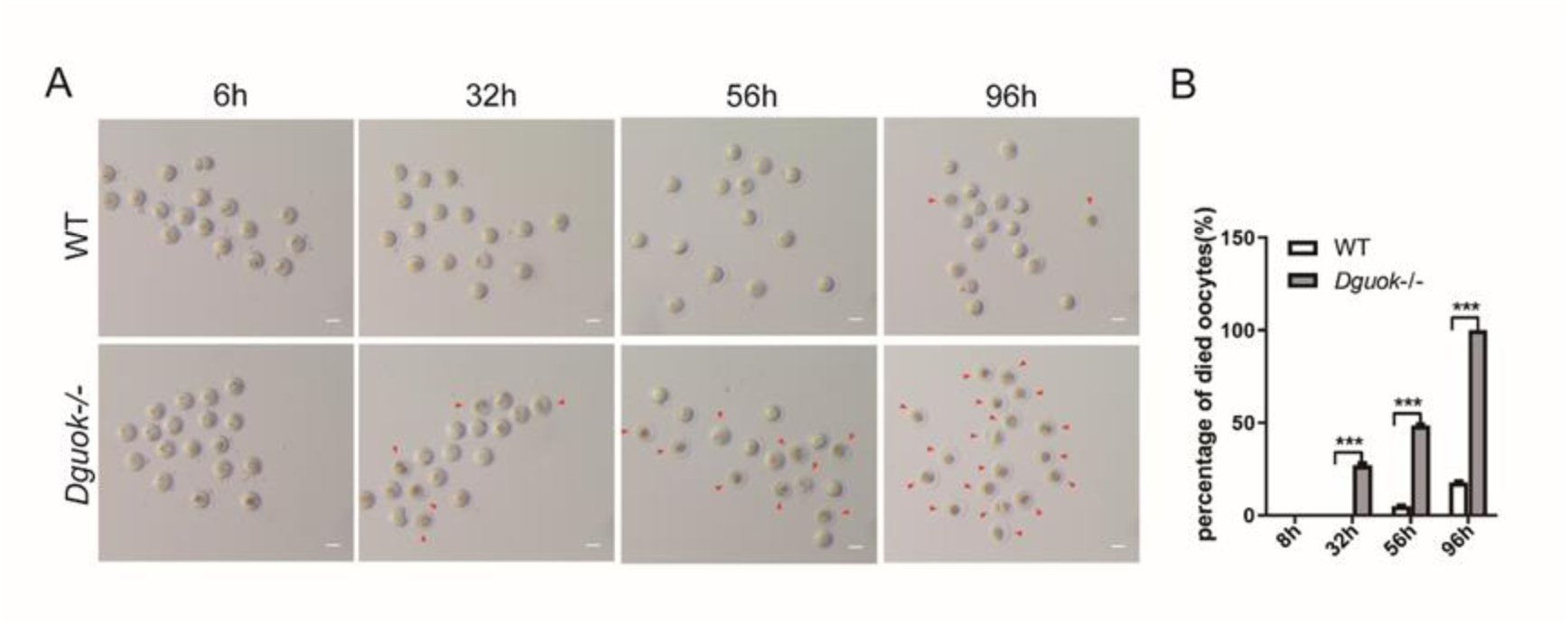
DGUOK deficiency significantly increased apoptosis of oocytes. A. Natural death profile of WT and *Dguok*^-/-^ mice Oocytes at 6h, 32h, 56h, and 96h, dead cells are indicated by red arrows; Scale bars: 50μm. B. Quantification of the died oocyte ratio from WT and *Dguok*^-/-^ mouse; n=3, ***p < 0.001.

## REFERENCE

1. Tamrakar SR, Bastakoti R. Determinants of Infertility in Couples. J Nepal Health Res Counc. 2019;17(1):85–9.

2. Maddirevula S, Coskun S, Alhassan S, Elnour A, Alsaif HS, Ibrahim N, et al. Female Infertility Caused by Mutations in the Oocyte-Specific Translational Repressor PATL2. Am J Hum Genet. 2017;101(4):603–8.

3. Feng R, Sang Q, Kuang Y, Sun X, Yan Z, Zhang S, et al. Mutations in TUBB8 and Human Oocyte Meiotic Arrest. N Engl J Med. 2016;374(3):223–32.

4. Zeng Y, Shi J, Xu S, Shi R, Wu T, Li H, et al. Bi-allelic mutations in MOS cause female infertility characterized by preimplantation embryonic arrest. Hum Reprod. 2022;37(3):612–20.

5. Huang HL, Lv C, Zhao YC, Li W, He XM, Li P, et al. Mutant ZP1 in familial infertility. N Engl J Med. 2014;370(13):1220–6.

6. De Vos M, Devroey P, Fauser BC. Primary ovarian insufficiency. Lancet. 2010;376(9744):911-21.

7. Golezar S, Ramezani Tehrani F, Khazaei S, Ebadi A, Keshavarz Z. The global prevalence of primary ovarian insufficiency and early menopause: a meta-analysis. Climacteric. 2019;22(4):403–11.

8. Ke H, Tang S, Guo T, Hou D, Jiao X, Li S, et al. Landscape of pathogenic mutations in premature ovarian insufficiency. Nat Med. 2023;29(2):483–92.

9. Jiao X, Ke H, Qin Y, Chen ZJ. Molecular Genetics of Premature Ovarian Insufficiency. Trends Endocrinol Metab. 2018;29(11):795–807.

10. Qin Y, Jiao X, Simpson JL, Chen ZJ. Genetics of primary ovarian insufficiency: new developments and opportunities. Hum Reprod Update. 2015;21(6):787–808.

11. Tucker EJ, Grover SR, Bachelot A, Touraine P, Sinclair AH. Premature Ovarian Insufficiency: New Perspectives on Genetic Cause and Phenotypic Spectrum. Endocr Rev. 2016;37(6):609–35.

12. Mehlmann LM. Stops and starts in mammalian oocytes: recent advances in understanding the regulation of meiotic arrest and oocyte maturation. Reproduction. 2005;130(6):791–9.

13. Eppig JJ. Coordination of nuclear and cytoplasmic oocyte maturation in eutherian mammals. Reprod Fertil Dev. 1996;8(4):485–9.

14. Vogt EJ, Meglicki M, Hartung KI, Borsuk E, Behr R. Importance of the pluripotency factor LIN28 in the mammalian nucleolus during early embryonic development. Development. 2012;139(24):4514–23.

15. Loutradis D, Kiapekou E, Zapanti E, Antsaklis A. Oocyte maturation in assisted reproductive techniques. Ann N Y Acad Sci. 2006;1092:235–46.

16. Ferreira EM, Vireque AA, Adona PR, Meirelles FV, Ferriani RA, Navarro PA. Cytoplasmic maturation of bovine oocytes: structural and biochemical modifications and acquisition of developmental competence. Theriogenology. 2009;71(5):836–48.

17. Guerin P, El Mouatassim S, Menezo Y. Oxidative stress and protection against reactive oxygen species in the pre-implantation embryo and its surroundings. Hum Reprod Update. 2001;7(2):175–89.

18. Van Blerkom J, Runner MN. Mitochondrial reorganization during resumption of arrested meiosis in the mouse oocyte. Am J Anat. 1984;171(3):335–55.

19. Van Blerkom J, Davis PW, Lee J. ATP content of human oocytes and developmental potential and outcome after in-vitro fertilization and embryo transfer. Hum Reprod. 1995;10(2):415–24.

20. Kennedy HJ, Pouli AE, Ainscow EK, Jouaville LS, Rizzuto R, Rutter GA. Glucose generates sub-plasma membrane ATP microdomains in single islet beta-cells. Potential role for strategically located mitochondria. J Biol Chem. 1999;274(19):13281–91.

21. Stojkovic M, Machado SA, Stojkovic P, Zakhartchenko V, Hutzler P, Gonçalves PB, et al. Mitochondrial distribution and adenosine triphosphate content of bovine oocytes before and after in vitro maturation: correlation with morphological criteria and developmental capacity after in vitro fertilization and culture. Biol Reprod. 2001;64(3):904–9.

22. Bhatti JS, Bhatti GK, Reddy PH. Mitochondrial dysfunction and oxidative stress in metabolic disorders - A step towards mitochondria based therapeutic strategies. Biochimica et biophysica acta Molecular basis of disease. 2017;1863(5):1066–77.

23. Bock FJ, Tait SWG. Mitochondria as multifaceted regulators of cell death. Nature reviews Molecular cell biology. 2020;21(2):85–100.

24. Sprenger HG, Langer T. The Good and the Bad of Mitochondrial Breakups. Trends in cell biology. 2019;29(11):888–900.

25. Hong X, Isern J, Campanario S, Perdiguero E, Ramírez-Pardo I, Segalés J, et al. Mitochondrial dynamics maintain muscle stem cell regenerative competence throughout adult life by regulating metabolism and mitophagy. Cell stem cell. 2022;29(9):1298–314.e10.

26. Eisner V, Picard M, Hajnóczky G. Mitochondrial dynamics in adaptive and maladaptive cellular stress responses. Nature cell biology. 2018;20(7):755–65.

27. Guillén-Samander A, Leonzino M, Hanna MG, Tang N, Shen H, De Camilli P. VPS13D bridges the ER to mitochondria and peroxisomes via Miro. The Journal of cell biology. 2021;220(5).

28. Santos TA, El Shourbagy S, St John JC. Mitochondrial content reflects oocyte variability and fertilization outcome. Fertil Steril. 2006;85(3):584–91.

29. Yu Y, Dumollard R, Rossbach A, Lai FA, Swann K. Redistribution of mitochondria leads to bursts of ATP production during spontaneous mouse oocyte maturation. J Cell Physiol. 2010;224(3):672–80.

30. Wassarman PM, Josefowicz WJ. Oocyte development in the mouse: an ultrastructural comparison of oocytes isolated at various stages of growth and meiotic competence. J Morphol. 1978;156(2):209–35.

31. Plagemann PG, Wohlhueter RM, Woffendin C. Nucleoside and nucleobase transport in animal cells. Biochim Biophys Acta. 1988;947(3):405–43.

32. Mandel H, Hartman C, Berkowitz D, Elpeleg ON, Manov I, Iancu TC. The hepatic mitochondrial DNA depletion syndrome: ultrastructural changes in liver biopsies. Hepatology. 2001;34(4 Pt 1):776-84.

33. Lin S, Huang C, Sun J, Bollt O, Wang X, Martine E, et al. The mitochondrial deoxyguanosine kinase is required for cancer cell stemness in lung adenocarcinoma. EMBO Mol Med. 2019;11(12):e10849.

34. Sun J, Hu Y, Bai W, Sun R, Shang R. Role and mechanism of mitochondrial deoxyguanosine kinase in lung cancer tumorigenesis. SCIENTIA SINICA Vitae. 2019;49(7):893–901.

35. Sang L, He YJ, Kang J, Ye H, Bai W, Luo XD, et al. Mitochondrial Deoxyguanosine Kinase Regulates NAD(+) Biogenesis Independent of Mitochondria Complex I Activity. Front Oncol. 2020;10:570656.

36. Zhou X, Curbo S, Zhao Q, Krishnan S, Kuiper R, Karlsson A. Severe mtDNA depletion and dependency on catabolic lipid metabolism in DGUOK knockout mice. Hum Mol Genet. 2019;28(17):2874–84.

37. Harris SE, Leese HJ, Gosden RG, Picton HM. Pyruvate and oxygen consumption throughout the growth and development of murine oocytes. Mol Reprod Dev. 2009;76(3):231–8.

38. Warzych E, Lipinska P. Energy metabolism of follicular environment during oocyte growth and maturation. J Reprod Dev. 2020;66(1):1–7.

39. Kidder BL. In vitro maturation and in vitro fertilization of mouse oocytes and preimplantation embryo culture. Methods in molecular biology (Clifton, NJ). 2014;1150:191–9.

40. Shadel GS, Clayton DA. Mitochondrial DNA maintenance in vertebrates. Annu Rev Biochem. 1997;66:409–35.

41. May-Panloup P, Boguenet M, Hachem HE, Bouet PE, Reynier P. Embryo and Its Mitochondria. Antioxidants (Basel). 2021;10(2).

42. Zhao S, Heng N, Wang H, Wang H, Zhang H, Gong J, et al. Mitofusins: from mitochondria to fertility. Cell Mol Life Sci. 2022;79(7):370.

43. Brevini TA, Vassena R, Francisci C, Gandolfi F. Role of adenosine triphosphate, active mitochondria, and microtubules in the acquisition of developmental competence of parthenogenetically activated pig oocytes. Biol Reprod. 2005;72(5):1218–23.

44. Wai T, Langer T. Mitochondrial Dynamics and Metabolic Regulation. Trends in endocrinology and metabolism: TEM. 2016;27(2):105–17.

45. Horbay R, Bilyy R. Mitochondrial dynamics during cell cycling. Apoptosis: an international journal on programmed cell death. 2016;21(12):1327–35.

46. Joaquim M, Escobar-Henriques M. Role of Mitofusins and Mitophagy in Life or Death Decisions. Frontiers in cell and developmental biology. 2020;8:572182.

47. Silva Ramos E, Larsson NG, Mourier A. Bioenergetic roles of mitochondrial fusion. Biochimica et biophysica acta. 2016;1857(8):1277–83.

48. Pagliuso A, Cossart P, Stavru F. The ever-growing complexity of the mitochondrial fission machinery. Cellular and molecular life sciences: CMLS. 2018;75(3):355–74.

49. Belli M, Palmerini MG, Bianchi S, Bernardi S, Khalili MA, Nottola SA, et al. Ultrastructure of mitochondria of human oocytes in different clinical conditions during assisted reproduction. Arch Biochem Biophys. 2021;703:108854.

50. Liu XM, Zhang YP, Ji SY, Li BT, Tian X, Li D, et al. Mitoguardin-1 and −2 promote maturation and the developmental potential of mouse oocytes by maintaining mitochondrial dynamics and functions. Oncotarget. 2016;7(2):1155–67.

51. Pan ZN, Pan MH, Sun MH, Li XH, Zhang Y, Sun SC. RAB7 GTPase regulates actin dynamics for DRP1-mediated mitochondria function and spindle migration in mouse oocyte meiosis. Faseb j. 2020;34(7):9615–27.

52. Moor RM, Dai Y, Lee C, Fulka J, Jr. Oocyte maturation and embryonic failure. Hum Reprod Update. 1998;4(3):223–36.

53. May-Panloup P, Boucret L, Chao de la Barca JM, Desquiret-Dumas V, Ferré-L’Hotellier V, Morinière C, et al. Ovarian ageing: the role of mitochondria in oocytes and follicles. Hum Reprod Update. 2016;22(6):725–43.

54. Yang L, Lin X, Tang H, Fan Y, Zeng S, Jia L, et al. Mitochondrial DNA mutation exacerbates female reproductive aging via impairment of the NADH/NAD(+) redox. Aging Cell. 2020;19(9):e13206.

55. Kristensen SG, Humaidan P, Coetzee K. Mitochondria and reproduction: possibilities for testing and treatment. Panminerva Med. 2019;61(1):82–96.

56. Qi J, Long Q, Yuan Y, Zhou Y, Zhang J, Ruan Z, et al. Mitochondrial DNA mutation affects the pluripotency of embryonic stem cells with metabolism modulation. Genome Instability & Disease. 2022;4(1):12–20.

57. Gershon E, Plaks V, Dekel N. Gap junctions in the ovary: expression, localization and function. Mol Cell Endocrinol. 2008;282(1-2):18–25.

58. Matsuda F, Inoue N, Manabe N, Ohkura S. Follicular growth and atresia in mammalian ovaries: regulation by survival and death of granulosa cells. J Reprod Dev. 2012;58(1):44–50.

